# Distress tolerance across self-report, behavioral and psychophysiological domains in women with eating disorders and healthy controls

**DOI:** 10.1101/170217

**Authors:** Angelina Yiu, Kara Christensen, Jean M. Arlt, Eunice Y. Chen

## Abstract

**Background and objectives:** The tendency to engage in impulsive behaviors when distressed is linked to engagement in disordered eating. The current study comprehensively examines emotional responses to a distress tolerance task by utilizing self-report, psychophysiological measures (respiratory sinus arrhythmia [RSA], skin conductance responses [SCRs] and tonic skin conductance levels [SCLs]), and behavioral measures (i.e., termination of task, latency to quit task).

**Methods:** 26 healthy controls (HCs) and a sample of treatment-seeking women with Bulimia Nervosa (BN), Binge Eating Disorder (BED) and Anorexia Nervosa (AN) (*N*=106) completed the Paced Auditory Serial Addition Task-Computerized (PASAT-C). Psychophysiological measurements were collected during baseline, PASAT-C, and recovery, then averaged for each time-period. Self-reported emotions were collected at baseline, post-PASAT-C and post-recovery.

**Results:** Overall, we found an effect of Time, with all participants reporting greater negative emotions, less happiness, lower RSA, more SCRs and higher tonic SCLs after completion of the PASAT-C relative to baseline. There were no differences in PASAT-C performance between groups. There was an effect of Group for negative emotions, with women with BN, BED and AN reporting overall higher levels of negative emotions relative to HCs. Furthermore, we found an effect of Group for greater urges to binge eat and lower RSA values among BED, relative to individuals with BN, AN and HCs.

**Limitations:** This study is cross-sectional and lacked an overweight healthy control group.

**Conclusion:** During the PASAT-C, individuals with EDs compared to HCs report higher levels of negative emotions, despite similar physiological and behavioral manifestations of distress.

---

Eating disorders (EDs) affect up to 4.64% of adults (Le Grange, Swanson, Crow, & Merikangas, 2012) and have significant medical and psychosocial outcomes (Baiano et al., 2014). Disordered eating, in general, has been conceptualized as a maladaptive response to alleviate distress, and thereby may negatively reinforce the disordered eating (Aldao, Nolen-Hoeksema, & Schweizer, 2010; Anestis, Smith, Fink, & Joiner, 2009; Corstorphine, Mountford, Tomlinson, Waller & Meyer, 2007; Fischer, Smith, & Cyders, 2008; Haynos & Fruzzetti, 2011; Leyro, Zvolensky, & Bernstein, 2010).

A number of studies use multiple methodologies to examine individuals’ with EDs experience of and abilities to withstand negative emotions and/or aversive states, termed distress tolerance. For example, it was found that self-report emotional distress tolerance, but not behavioral and physical forms of distress tolerance, was negatively associated with symptoms of bulimia nervosa (BN) (Anestis et al., 2012). Another study found that BN did not differ from healthy controls (HCs) on heat and cold pain thresholds after a stress induction task (Schmahl et al., 2010). While these studies expand our understanding of the utility of self-report and behavioral measures of distress tolerance in EDs, our understanding of the course of affective responding in EDs before, during, and after a stressor is still limited (Anestis et al., 2007; Anestis, Smith, et al., 2009; Claes, Vandereycken, & Vertommen, 2005; Peterson & Fischer, 2012; Wenzel, Weinstock, Vander Wal, & Weaver, 2014; Wu et al., 2013).

To gain a comprehensive understanding of affective response when engaged in distress tolerance in EDs, multiple methods of assessment are needed (Gross, 2013); however, there are few studies that utilize this approach, therefore any relationship between the psychophysiological components of emotional responding and the behavioral components in EDs remains unclear (Gross, 2013; Lang, Greenwald, Bradley, & Hamm, 1993).

The psychophysiological component of emotional response is comprised of parasympathetic and sympathetic nervous system activity. Parasympathetic nervous system activity can be indexed by respiratory sinus arrhythmia (RSA), which consists of heart rate variability (HRV) in conjunction with respiration. Neuroimaging studies show that there is a positive relationship between RSA and increased activation in neural structures implicated in emotion processing and physiological aspects of emotional responses (e.g., right pregenual anterior cingulate, right subgenual anterior cingulate and right rostral medial prefrontal cortex and the left sublenticular extended amygdala or ventral striatum) (Thayer, Åhs, Fredrikson, Sollers, & Wager, 2012). Reduced resting RSA is theorized as a risk factor for psychopathology (Crowell, Beauchaine, & Linehan, 2009), although systematic reviews and meta-analysis suggest that higher resting HRV is associated with both BN (Peschel et al., 2016) and anorexia nervosa (AN) (Mazurak, Enck, Muth, Teufel, & Zipfel, 2011) compared to controls. Low resting RSA, decreased RSA when emotionally aroused, and a slow return to baseline after emotional arousal are associated with greater symptoms of psychopathology (Beauchaine, 2015).

Experimental work examining RSA response to stressors among individuals with EDs has produced mixed findings. Two separate studies found that women with BED and obesity (Friederich et al., 2006) and women with BN (Messerli-Bürgy, Engesser, Lemmenmeier, Steptoe, & Laederach-Hofmann, 2010) showed decreased RSA levels with a slow return to baseline following a psychological stress induction task. However, there is also evidence to suggest that RSA levels among women with BED and obesity did not change after psychological stress was induced (Messerli-Bürgy et al., 2010) and conversely women without EDs who are obese showed a return to baseline RSA levels after stress exposure (Messerli-Bürgy et al., 2010). Additional research suggests that decreased RSA after psychological stress is associated with engagement in clinically significant levels of binge eating, rather than weight status (Udo et al., 2014).

The sympathetic nervous system response can be indexed by skin conductance response, which measures the time it takes for a current to pass through the skin (Boucsein et al., 2012). Higher skin conductance responses (SCRs) are associated with greater emotional arousal (Kreibig, 2010; Lang et al., 1993). In contrast to the mixed relationship between RSA and EDs, evidence suggests that skin conductance levels (SCL) are similar between women with BN and BED in response to different stressors (Hilbert, Vögele, Tuschen-Caffier, & Hartmann, 2011) and between women with BN, women with self-reported restrained eating or HCs (Tuschen-Caffier & Vögele, 1999). Taken together, there is some evidence to suggest that there is a relationship between EDs and decreased RSA when psychological stress is induced, and little evidence pointing to a relationship between EDs and sympathetic responses as indexed through skin conductance.

Thus far, the evidence base for the psychophysiological component of emotional response while distressed across the range of EDs has been mixed, limited by small sample sizes, and there is a lack of research utilizing multiple ED diagnoses. The current study extends past research (Leehr et al., 2015; Naumann, Tuschen-Caffier, Voderholzer, Caffier, & Svaldi, 2015; Svaldi, Griepenstroh, Tuschen-Caffier, & Ehring, 2012; Svaldi, Tuschen-Caffier, Lackner, Zimmermann, & Naumann, 2012) through the multi-modal assessment of emotional responses across three common ED diagnoses and HC participants to a commonly used behavioral distress tolerance task. The Paced Auditory Serial Addition Task-Computerized (PASAT-C) (Lejuez, Kahler, & Brown, 2003) was used as a behavioral measure of distress tolerance (Feldner, Leen-Feldner, Zvolensky, & Lejuez, 2006; Gratz, Rosenthal, Tull, Lejuez, & Gunderson, 2006) and self-reported emotions and psychophysiological measures of arousal were measured in response to the PASAT-C. As individuals with EDs have been argued to have, “shared, but distinctive, clinical features […] maintained by similar psychopathological processes,” utilizing a transdiagnostic approach that includes three ED groups may be most useful in identifying the proposed “similar psychopathological processes” (Fairburn, Cooper, & Shafran, 2003)

We predicted a main effect of time, such that self-reported negative emotions would be greater, happiness would be lower, and urges to binge eat would be greater after the PASAT-C, in comparison to baseline and at recovery. Furthermore, we expected that diagnostic group would moderate this effect. Specifically, we expected that HCs would demonstrate less significant changes in negative emotions, happiness, and urges to binge eat during the PASAT-C, compared to individuals with BN, BED or AN. As the PASAT-C is used as a demanding behavioral distress tolerance measure (Gratz et al., 2006; Sauer & Baer, 2012), we expected that individuals with BN, BED and AN would be more likely to prematurely terminate the PASAT-C and exhibit shorter latency to terminate the PASAT-C than HCs. Given the proposed role of negative affect in EDs during distress (Anestis, Peterson, et al., 2009; Anestis et al., 2007; Peterson & Fischer, 2012), we expected that RSA values would be lower and SCRs and tonic SCL values would be higher during the PASAT-C in ED groups compared to HC, relative to baseline and recovery.

## Methods

### Participants

Participants were self-referred from the community using flyers, referred from local eating disorder clinics and student health and counseling services, and recruited in a University-based outpatient eating disorder program as part of several clinical trials. Participants were invited to participate in the study if, after undergoing a clinical interview (described below) they met for a diagnosis of BN, BED or AN (typical or atypical), without current drug or alcohol dependence or symptoms of psychosis. HCs were self-referred from the community, student health and counseling services, and the University Hospital, and were eligible to participate if, after undergoing the same clinical interview as EDs, they did not meet diagnosis for current psychopathology, and did not have a history of serious mental illness.

The sample consisted of 106 females with an ED and 26 female HCs with no history of psychiatric disorders from the community. Of the participants with an ED, 34.90% of the sample was diagnosed with BN (*n*=37), 51.89% with BED (*n*=55), and 13.21% with typical or atypical AN (*n*=14). The sample identified predominantly as non-Hispanic (91.70%) and were 64.4% Caucasian, 13.60% African-American, 4.50% Asian, and 9.1% of mixed race. The mean age of participants was 35.06 years (*SD* = 11.47), and the mean body mass index (BMI) was 29.55 kg/m^2^ (*SD* = 9.50).

### Measures

#### Psychiatric symptoms

The Structured Clinical Interview for Diagnostic and Statistical Manual of Mental Disorders – IV-Text Revision (DSM – IV – TR) Axis I Disorders (SCID – I) (First, Spitzer, Gibbon, & Williams, 2002) was used to assess for psychiatric symptoms. The SCID I was used to confirm ED diagnoses with the EDE-16.0 and to assess for exclusionary psychiatric symptoms. The SCID I is considered the gold-standard measure for psychiatric diagnoses (Lobbestael, Leurgans & Arntz, 2011).

#### Eating disorder behaviors

The Eating Disorder Examination 16.0 (EDE) (Fairburn, Cooper, & O’Connor, 2008) is an investigator-based interview that assesses DSM– IV-TR (American Psychiatric Association, 2000) ED symptoms. The EDE – 16.0 was used to diagnosis AN, BN and BED and generate subscales that tap into ED behaviors and cognitions: Eating Concern, Shape Concern, Weight Concern, Dietary Restraint, as well as a Global score. The EDE – 16.0 was also used to assess for the number of binge eating and compensatory episodes in the prior 3-month period. The EDE – 16.0 has good discriminant validity, such that individuals with AN, BN and BED are distinguished from controls (Cooper, Cooper & Fairburn, 1989; Wilson & Smith, 1989; Wilfley, Schwartz, Spurrell & Fairburn, 1999). In the current study, the EDE total score demonstrated an internal consistency of α = .94.

#### Current subjective emotional state

A visual analogue scale (VAS) with an abbreviated Positive and Negative Affect State (PANAS) (Watson, Clark, & Tellegen, 1988) measured current subjective emotional state prior to the baseline, after the PASAT-C, and after recovery from the PASAT-C (see Figure 1). Self-report negative and positive affective adjectives that assessed anxiety, fear, frustration, happiness, sadness, and tension were scored on a 100-point Likert scale. A single question to assess urges to binge eat was added due to its relevance in the current ED sample. Higher scores indicated greater intensity of the response. In the present study, we created a composite for negative emotions using an average of scores from anxiety, fear, frustration, sadness and tension. Urge to binge eat and happiness were assessed separately. The original PANAS shows good convergent validity, with correlations ranging from .51 to .74 with the Beck Depression Inventory, the State-Trait Anxiety Inventory State Anxiety Scale and Hopkins Symptom Checklist (Watson, Clark & Tellegen, 1988). In the present study, internal consistency of the negative emotion composite score at each time point ranged from α = .84 to .90.

**Figure 1.**
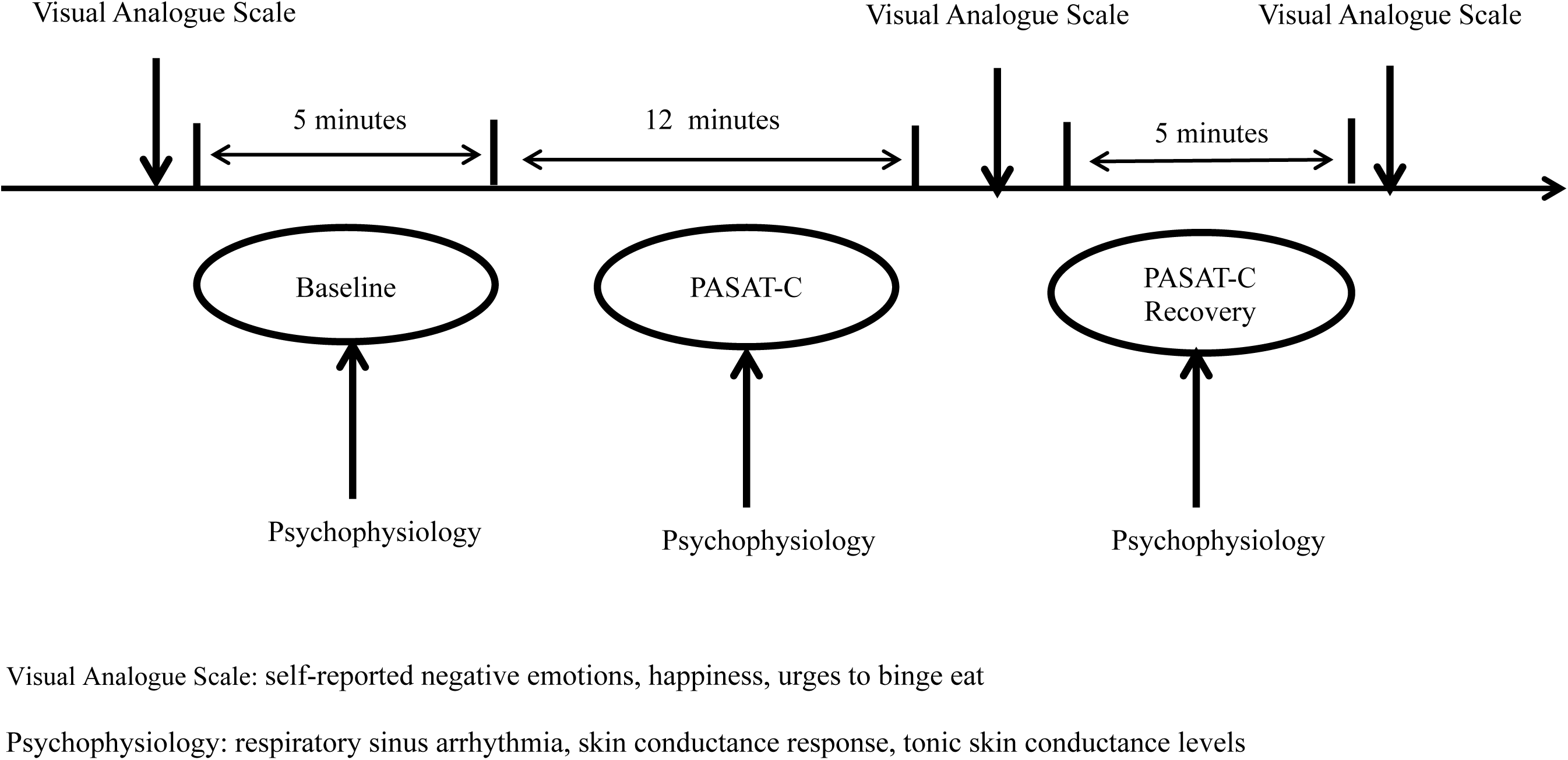
Schematic of the procedure and time periods assessed.

#### Psychophysiological measures

Psychophysiological measures included respiratory sinus arrhythmia (RSA) as a measure of parasympathetic activity (Grossman & Taylor, 2007; Thayer et al., 2012) and skin conductance responses (SCRs) and tonic skin conductance levels (tonic SCL) as measures of sympathetic activity (Boucsein, 2011). Psychophysiological measures were assessed during the baseline, during the PASAT-C and during the recovery period. Electrocardiogram (ECG) data were collected from two Biopac Ag-AgCl spot electrodes placed on the bottom of the palm of the non-dominant hand, using a modified Lead II configuration (Boucsein, 2011). We derived RSA by using a band pass filter on the ECG signal and spectral analysis to extract the high-frequency component (> .15 hz) of heart rate variability. Skin conductance was recorded from two electrodes attached to the palm of the non-dominant hand. Increases in SCRs have been associated with negative affect following exposure to emotional stimuli, even when cardiac measures did not change significantly (Boucsein et al., 2012; Salters-Pedneault, Gentes, & Roemer, 2007). Tonic SCL referred to the level of skin conductance per data collection period.

**Diagnostic and Laboratory Procedures**.

The present study was reviewed and approved by the university institutional review board and took place over two separate testing sessions, with each session taking 2-4 hours. During the first session, eligibility screening and informed consent were completed, followed by an assessment of psychiatric and ED symptoms by Masters-level clinicians using the measures described below. Individual participant diagnoses were agreed upon and finalized with the licensed clinical psychologist supervisor at weekly best-estimate meetings (Klein, Ouimette, Kelly, Ferro, & Riso, 1994; Kosten & Rounsaville, 1992), where all structured interview data was presented. Height and weight were measured to calculate BMI. During the second session, participants completed the laboratory procedures for the PASAT-C, described below. Diagnostic and psychophysiological assessments occurred prior to outpatient treatment for individuals with EDs.

#### Behavioral distress tolerance task

The Paced Auditory Serial Addition Task-Computerized (PASAT-C) (Lejuez et al., 2003) is a behavioral distress tolerance task that has been shown to induce negative affect (Daughters, Lejuez, Kahler, Strong, & Brown, 2005; Feldner et al., 2006; Holdwick & Wingenfeld, 1999; Schloss & Haaga, 2011; Eichen, Chen, Boutelle & McCloskey, 2017) and short-term anxiety, frustration, and irritability (Gratz et al., 2006; Lejuez et al., 2003). The PASAT-C is a computer-based task that requires the participant to add a visually presented digit to the previous visually presented digit. Explosion sounds followed incorrect answers or when the participant failed to respond quickly enough. The PASAT-C was presented for a maximum of twelve minutes, consisting of four levels that lasted for three minutes each. With each successive level of the PASAT-C, the latency between trials was decreased and negative feedback of “*go faster*,” *“do better*,” and “*go faster, do better*” was added. By the fourth level (PASAT-C Level 4), there was a one second latency of trials with negative feedback. Participants were provided written instructions at the beginning of the task that they had the option to quit the task after completion of Level 2. If participants terminated the task early, they went on to complete the recovery period, detailed below. Distress tolerance on the PASAT-C was behaviorally operationalized dichotomously and continuously as termination of the task when given the option and latency to quit, respectively.

#### Psychophysiology Capture Procedure

Psychophysiological measures were collected during baseline, PASAT-C, and the recovery period. Average RSA, SCR and tonic SCL values were computed from the 5-minute baseline, up to 12 minutes of the PASAT-C, and the 5-minute recovery period, respectively. Please see Figure 1 for a schematic of the procedure. Once sensors were attached, participants completed a baseline VAS and were asked to sit quietly without moving for a 5-minute baseline period (Laborde, Mosley & Thayer, 2017). Following this, participants were instructed to begin the PASAT-C. Upon completion of the PASAT-C, participants completed a second VAS and were asked to sit quietly for an additional 5-minutes without moving for a recovery period (Laborde et al., 2017). Participants completed a final VAS rating after the recovery period.

### Statistical Treatment

Chi-square tests were conducted to assess for group differences in all demographic variables and medical and psychiatric comorbidities, with the exception of age, which was assessed using an ANOVA. Variables related to presence of medication use (e.g., stimulant medication, anxiolytics) and medical co-morbidities were included as covariates in the analyses if there were significant differences between ED groups (BN, BED, AN) due to their effects on psychophysiological responses (Grossman, Stemmler, & Meinhardt, 1990; Masi, Hawkley, Rickett, & Cacioppo, 2007). As it was expected that HCs would exhibit significantly less medical co-morbidities and medication use relative to the ED groups (Hudson, Hiripi, Pope, & Kessler, 2007; Johnson, Spitzer, & Williams, 2001; Kessler et al., 2013; Padierna, Quintana, Arostegui, Gonzalez, & Horcajo, 2000), medical co-morbidities and medication use were only included as covariates if there were significant differences between BN, BED and AN.

For analyses involving self-reported emotions and psychophysiological measures, independent variables were Time (baseline, PASAT-C, recovery) and Group (BN, BED, AN and HC). A repeated measures ANOVA was chosen as it allows for individual differences in baseline scores to be accounted (Field, 2013). Six repeated measures Time x Group ANOVAs were conducted with the self-reported negative emotion composite score, happiness, urge to binge eat, RSA, SCRs and Tonic SCL as dependent variables. Planned simple effects analyses were conducted to probe significant interactions. To examine group differences in PASAT-C completion vs. non-completion, a chi square test was conducted. To examine group differences in PASAT-C latency to quit, a univariate ANOVA was conducted. A power analysis indicated that we were powered to detect medium to large interaction effects (*f* = .25 to .40) with a power of .95 (Faul, Erdfelder, Lang, & Buchner, 2007). Partial *η*^2^ was used to report effect sizes for repeated measures ANOVAs and ANOVAs, with the following cut-off conventions: small (.01), medium (.06) and large (.14) (Cohen, 1988).

## Results

### Preliminary Analyses

As expected, there were significant group differences on presence of a lifetime mood disorder (*p* < .001), anxiety disorder (*p* < .001), medical co-morbidities (*p* = .006) and medication use (*p* = .002), which were driven by the lack of psychiatric and medical co-morbidities among HCs. There were no significant differences between BN, BED and AN on presence of a lifetime mood disorder, medical co-morbidities or medication use, although AN were significantly more likely to have an anxiety disorder relative to BN (*p* < .05). Due to the lack of significant systematic differences in psychiatric and medical co-morbidities between BN, BED and AN, these variables were not included as covariates. Please see Table 1 for a description of the severity of ED symptoms, demographic information and clinical characteristics of the sample.

**Table 1.**
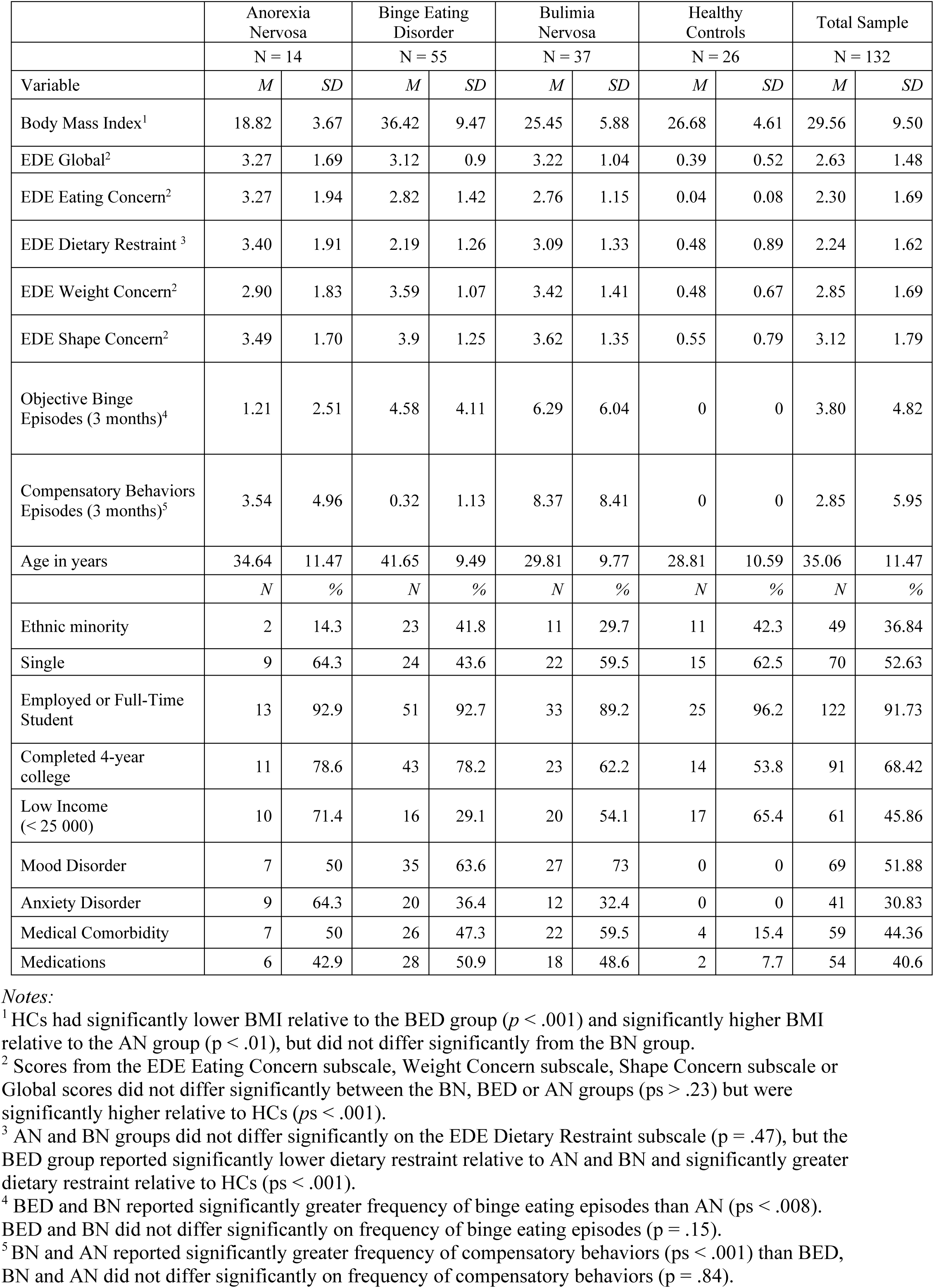
Severity of eating disorder symptoms, demographic information and clinical characteristics of participants at baseline.

**Table 2.** Self-reported negative emotions, happiness and urges to binge eat at baseline, during the Paced Auditory Serial Addition Task – Computerized (PASAT-C), and during recovery from PASAT-C among women with Bulimia Nervosa, Binge Eating Disorder, Anorexia Nervosa and Healthy Controls.

|  | Baseline<br>*M (SD)* | PASAT-C<br>*M (SD)* | Recovery<br>*M (SD)* |
| --- | --- | --- | --- |
| Negative Emotions |  |  |  |
| Anorexia Nervosa | 48.99 (22.51) | 60.80 (26.58) | 47.53 (21.66) |
| Binge Eating Disorder | 39.40 (22.82) | 57.24 (20.20) | 30.55 (18.25) |
| Bulimia Nervosa | 38.90 (22.83) | 58.38 (20.22) | 35.90 (20.33) |
| Healthy Control | 8.91 (13.18) | 37.07 (24.47) | 15.40 (16.38) |
| Happiness |  |  |  |
| Anorexia Nervosa | 37.56 (25.16) | 22.47 (26.08) | 17.68 (24.51) |
| Binge Eating Disorder | 40.85 (18.05) | 23.52 (25.07) | 29.20 (21.58) |
| Bulimia Nervosa | 43.98 (18.74) | 21.03 (21.58) | 29.50 (23.21) |
| Healthy Control | 59.41 (19.40) | 28.51 (28.71) | 39.42 (28.14) |
| Urge to Binge |  |  |  |
| Anorexia Nervosa | 21.70 (29.03) | 11.42 (25.86) | 20.41 (33.67) |
| Binge Eating Disorder | 46.87 (27.10) | 42.41 (36.48) | 44.21 (32.50) |
| Bulimia Nervosa | 33.15 (28.48) | 31.84 (36.86) | 34.77 (32.55) |
| Healthy Control | .77 (1.97) | 2.98 (11.41) | .34 (.71) |

### Main Analyses^1^

#### Hypothesis 1: Self-reported emotions

Overall, there was a main effect of Time, such that participants reported higher negative emotions, *F*(2, 107)= 47.61, *p* < .001, η^2^=.31, and lower happiness, *F*(2, 107)= 39.10, *p* < .001, η^2^=.27, after completion of the PASAT-C, in comparison to baseline and recovery (*ps* < .05). There was a main effect of Group for negative emotions, F_(3,107)_ = 12.09, *p* < .001, η^2^=.25, but not happiness (*p* = .07). Specifically, HC participants reported lower negative emotions (*ps* < .001) in comparison to individuals with BN, BED or AN across all measurement periods. There were no Time x Group interactions (*ps* = .12 - .27) for negative emotions or happiness.

For urges to engage in binge eating, there was a main effect of Group, *F*(3, 107)= 16.51, *p* < .001, η ^2^ = .32, such that across all measurement periods individuals with BED demonstrated significantly greater urges to binge eat compared to BN, AN and HCs (*ps* < .04), individuals with BN demonstrated significantly greater urges to binge eat compared to HCs (*p* < .001), and individuals with AN did not differ significantly from BN or HC (*p*s > .06). There was no effect of Time (*p* = .56) or an interaction between Time x Group (*p* = .93), but individuals with AN exhibited a trend for decreased urge to binge eat after completion of the PASAT-C. See Table 3 for a summary of findings.

**Table 3.** Completion time and latency to quit on the Paced Auditory Serial Addition Task – Computerized (PASAT-C) among women with Bulimia Nervosa, Binge Eating Disorder, Anorexia Nervosa and Healthy Controls.

|  | PASAT-C<br>Completion<br>N (%) | PASAT-C<br>Latency to Quit (seconds)<br>M (SD) |
| --- | --- | --- |
| Anorexia Nervosa | 10 (71.4%) | 678.64 (103.62) |
| Binge Eating Disorder | 38 (69.1%) | 683.84 (89.32) |
| Bulimia Nervosa | 23 (62.2%) | 643.84 (128.05) |
| Healthy Control | 22 (84.6%) | 690.85 (110.35) |

#### Hypothesis 2: PASAT-C termination and latency to quit

A chi square test indicated that there were no differences between groups on PASAT-C completion versus non-completion, *χ*^2^(3)= 3.85, *p* = .28. A univariate ANOVA suggested that there were no differences between. groups on PASAT-C latency to quit, *F*(3,129) = 1.39, *p* = .25, η^2^ = .03. See Table 4 for means and standard deviations.

**Table 4.** Respiratory sinus arrhythmia, skin conductance responses and tonic skin conductance levels for Bulimia Nervosa, Binge Eating Disorder, Anorexia Nervosa and Healthy Controls at baseline, during PASAT-C, and during recovery from PASAT-C.

|  | Baseline<br>*M* (SD) | PASAT-C<br>*M* (SD) | Recovery<br>*M* (SD) |
| --- | --- | --- | --- |
| Respiratory sinus arrhythmia (in ms<sup>2</sup>) |  |  |  |
| Anorexia Nervosa | 5.81 (1.20) | 5.69 (1.16) | 6.15 (1.17) |
| Binge Eating Disorder | 5.25 (1.36) | 5.16 (1.39) | 5.40 (1.32) |
| Bulimia Nervosa | 6.25 (1.16) | 6.05 (1.06) | 6.32 (1.09) |
| Healthy Control | 6.21 (1.36) | 6.06 (1.30) | 6.38 (1.27) |
| Skin conductance response |  |  |  |
| Anorexia Nervosa | 1.80 (2.42) | 3.96 (4.15) | 3.91 (2.69) |
| Binge Eating Disorder | 2.13 (1.90) | 3.78 (3.02) | 3.88 (2.75) |
| Bulimia Nervosa | 1.24 (1.70) | 3.54 (3.45) | 2.77 (2.39) |
| Healthy Control | 1.83 (1.72) | 2.90 (2.54) | 3.94 (2.94) |
| Tonic skin conductance levels (in μs) | | | |
| Anorexia Nervosa | 2.19 (2.11) | 6.38 (6.58) | 6.72 (5.67) |
| Binge Eating Disorder | 2.94 (2.27) | 5.14 (4.06) | 5.06 (4.05) |
| Bulimia Nervosa | 2.35 (3.89) | 5.28 (6.51) | 4.98 (6.40) |
| Healthy Control | 2.40 (3.11) | 5.37 (6.04) | 5.77 (6.76) |

#### Hypothesis 3: Psychophysiological measures

There was a significant effect of Time for RSA values, *F*(2,128) = 11.87, *p* < .001, η^2^= .09, such that all participants exhibited lowered RSA during the PASAT-C, in comparison to baseline (*p* = .05) and recovery (*p* < .001). There was a significant effect of Group, *F*(3,128) = 6.10, *p* = .001, η^2^= .13, such that individuals with BED exhibited lowered RSA levels in comparison to individuals with BN or HCs (*ps* ≤ .006) but not AN (*p* = .325) across all measurement periods. There was no Time x Group interaction for RSA values.

There was a significant effect of Time for Tonic SCL values, *F*(2,109) = 50.57, *p* < .001, η^2^= .32, and SCR values, *F*(2,107) = 32.85, *p* < .001, η^2^= .24, such that all participants exhibited greater Tonic SCL and SCR values during the PASAT-C in comparison to baseline (*ps* < .001), but not recovery (*ps* =.45 - .75). There was no effect of Group or Time x Group interaction for Tonic SCL or SCR values. See Table 5 for means and standard deviations.

## Discussion

The current study extends the current literature on affective response in EDs by integrating self-report, behavioral and psychophysiological measures (RSA, SCRs and tonic SCLs) of response to a distress tolerance task (Daughters et al., 2005; Feldner et al., 2006; Holdwick & Wingenfeld, 1999; Schloss & Haaga, 2011). Despite similar psychophysiological responding (RSA, SCRs and tonic SCL) to the PASAT-C and PASAT-C termination and latency to quit across all groups, individuals with BN, BED and AN reported greater overall negative emotions and lower overall happiness compared to HCs. The discrepancy between psychophysiological responding and PASAT-C termination and latency to quit with self-report of emotional experience suggests that individuals with EDs experience disproportionately greater subjective distress relative to HCs when presented with the same stimuli. Overall, this may suggest that while physiological and behavioral experience of distress manifests similarly across groups, it is the interpretation of experiences during both baseline and when distressed that best distinguish between groups with EDs and HCs.

Although HCs reported significantly lower negative emotions and greater happiness relative to all other groups, individuals with BED were differentiated from other ED groups and HCs by exhibiting significantly lower RSA values without concomitant differences from other groups in Tonic SCL or SCR values. Research has demonstrated an association between BMI and RSA and suggests that cardiovascular autonomic function changes as a function of weight loss (Karason, Mølgaard, Wikstrand, & Sjöström, 1999; Rissanen, Franssila-Kallunki, & Rissanen, 2001). Consistent with past research (Dingemans & van Furth, 2012; Hudson et al., 2007), HCs had significantly lower BMI relative to the BED group and significantly higher BMI relative to the AN group, but did not differ significantly from the BN group; however, BMI was not controlled for in the current study, as 92% of participants diagnosed with BED were overweight or obese, likely due to the cumulative effects of objective binge episodes on weight gain over time (Hudson et al., 2010). Therefore, participants with BED are “doubly diagnosed” with BED and obesity, making it difficult to dissociate diagnosis and weight status. Future studies may wish to dissociate diagnosis and weight status.

Contrary to expectations, we did not find a Time (baseline, PASAT-C and recovery) by Group interaction in self-reported urge to binge eat. However, there were Group differences in urge to binge eat, such that individuals with BED and BN reported overall greater urges to binge eat across all time-points in comparison to individuals with AN or HC. Perhaps individuals who engage in habitual binge eating experience consistent urges to engage in binge eating, but only act on this urge when they experience negative affect (Hilbert & Tuschen-Caffier, 2007) or are under stress. As the current study only assessed urges to binge eat, future research may seek to explore factors that influence whether one will or will not act upon urges to binge eat when experiencing negative affect (Haedt-Matt & Keel, 2011; Hilbert & Tuschen-Caffier, 2007).

The PASAT-C achieved the desired result of increasing distress as evidenced by increased negative emotions, decreased happiness, decreased RSA values and increased SCRs and tonic SCLs across the sample, without differences in termination or latency to quit. This suggests that regardless of baseline level of negative emotions or diagnostic group, the PASAT-C was distress inducing, which is consistent with past research utilizing samples with clinically significant psychopathology (Gratz et al., 2006). The success of the PASAT-C to induce distress may have produced a ceiling effect on performance, such that all participants were equally sensitive to its effects regardless of diagnosis. However, the lack of significant group differences on the PASAT-C is counter to another study that found significantly shorter latency to quit the PASAT-C among undergraduate students who endorsed binge eating behaviors relative to controls (Eichen et al., 2017). The difference in findings may be due to sampling differences between the two studies. The current study recruited treatment-seeking patients with a range of EDs; the other study recruited non-treatment seeking college students.

## Future Directions

The inclusion of multiple ED diagnostic groups can be considered a strength and a limitation of the current study. Comparisons were possible between ED diagnoses and HCs, however the smaller group of individuals with AN may have hindered detecting differences between that group and others. For example, individuals with AN exhibited a trend for decreased urges to binge eat after completion of the PASAT-C in comparison to baseline, which is in contrast to individuals with BED and BN who exhibited similar urges to binge eat at baseline and after the PASAT-C. This may reflect an increased desire for control over eating when experiencing negative affect among individuals with AN (Fairburn, Shafran, & Cooper, 1999). A larger sample of individuals with AN may clarify the urge to binge eat in response to stress among different ED diagnoses. Furthermore, as the PASAT-C generated similar levels of negative affect across the entire sample, future research could utilize emotion inductions that may be more salient to EDs, such as cues involving body shape/weight concerns or food. Such disorder-related stimuli may produce differential results between groups with EDs and HCs and offer increased insight into the negative affect states that are posited to maintenance disordered eating behaviors. For example, participants could complete behavioral approach/avoidance tasks that involve examining the body in a mirror or observing images of high-caloric food. In terms of behavioral assessments, future research could utilize distress tolerance measures that more closely map onto eating disorder concerns, such as sampling foods.

Finally, the current study’s group of individuals with BED primarily consisted of individuals who are also overweight. Although individuals with BED are at a higher risk for obesity (e.g., de Zwaan, 2001), the medical condition of obesity is not synonymous with the psychological condition of BED (e.g., Klatzkin, Gaffney, Cyrus, Bigus, & Brownley, 2015). There are multiple physiological, cognitive, and psychological symptoms associated with obesity and there is a risk of falsely conflating these symptoms with BED. Future studies may seek to disentangle BED and obesity by examining whether similar patterns of findings are found for individuals with BED who are of normal weight, or for individuals without EDs who are of normal weight and overweight status.

The current study is one of the first to examine emotional responding across individuals with EDs and HCs in response to a task that induces distress. It provides a step towards enhancing our understanding of the similarities and differences in emotional responding across ED diagnoses using multiple measures of emotional responding. As previously suggested, our finding that individuals with BED exhibit overall reduced levels of RSA responding and experience consistent urges to binge eat over time requires further study before treatment implications can be made. The finding that EDs are associated with overall greater subjective distress without deleterious effects on the PASAT-C termination or latency to quit relative to HCs has treatment implications for the function of disordered eating to decrease distress (Corstorphine et al., 2007). Overall, performance on the PASAT-C was comparable across groups, suggesting there are not differences in the cognitive functioning required for the task, thus clinical intervention for EDs may prioritize managing strong, negative emotion. Individuals with EDs may benefit from treatments that focus on fostering greater acceptance of one’s emotional experiences, such as Dialectical Behavior Therapy (Chen et al., 2015; Safer, Robinson, & Jo, 2010) or Acceptance and Commitment Therapy (Berman, Boutelle, & Crow, 2009; Juarascio et al., 2013; Juarascio, Forman, & Herbert, 2010). Our study supports the proposed model that negative emotions may perpetuate and maintain EDs, with the caveat that the experience of distress may manifest physiologically and behaviorally similarly to HCs.

## Footnotes

1 Analyses were re-run with BMI as a covariate, with some differences in findings. Specifically, there was no longer an effect of Time for negative emotions (*p* = .70) or RSA (*p* = .83). All other findings remained the same.

